# Plant neopolyploidy and genetic background differentiates the microbiome of duckweed across a variety of natural freshwater sources

**DOI:** 10.1101/2023.04.29.538806

**Authors:** Thomas J. Anneberg, Martin M. Turcotte, Tia-Lynn Ashman

**Affiliations:** Department of Biological Sciences, University of Pittsburgh, Pittsburgh, PA, USA 15260

**Keywords:** Araceae, plant-microbe symbiosis, microbial community assembly, synthetic polyploidy, whole genome duplication

## Abstract

Whole genome duplication has long been appreciated for its role in driving phenotypic novelty in plants, often altering the way organisms interface with the abiotic environment. Only recently, however, have we begun to investigate how polyploidy influences interactions of plants with other species, despite the biotic niche being predicted as one of the main determinants of polyploid establishment. Nevertheless, we lack critical information about how polyploidy affects the diversity and composition of the microbial taxa that colonize plants, and whether this is genotype-dependent and repeatable across natural environments. This information is a critical first step toward understanding whether the microbiome contributes to polyploid establishment. We thus tested the immediate effect of polyploidy on the diversity and composition of the bacterial microbiome of the aquatic plant *Spirodela polyrhiza* using four pairs of diploids and synthetic autotetraploids. Under controlled conditions, axenic plants were inoculated with pond waters collected from 10 field sites across a broad environmental gradient. Autotetraploids hosted 4-11 % greater bacterial taxonomic and phylogenetic diversity than their diploid progenitors. Polyploidy, along with its interactions with the inoculum source and genetic lineage, collectively explained 7 % of the total variation in microbiome composition. Furthermore, polyploidy broadened the core microbiome, with autotetraploids having 15 unique bacterial taxa in addition to the 55 they shared with diploids. Our results show that whole genome duplication directly leads to novelty in plant microbiome and importantly, that the effect is dependent on the genetic ancestry of the polyploid and generalizable over many environmental contexts.

## Introduction

Whole genome duplication (“polyploidy”), which causes organisms to have greater than two sets of each chromosome, is a major force in ecology and evolution, especially for plants (Fox, Soltis, Soltis, Ashman, & Van de Peer, 2020; Hao et al., 2022; Rice et al., 2019; Wood et al., 2009). A key consequence of polyploidy is that it causes phenotypic novelty at all levels of biological complexity, from subcellular traits (e.g., nucleus size) to entire populations (e.g., population growth rate; (Anneberg et al., 2023; Doyle & Coate, 2019), and this has long-been appreciated to lead to novelty in the ecology of organisms (Soltis, Visger, & Soltis, 2014). Yet our understanding of the factors that contribute to the success of polyploids is dominated by evidence of their interactions with the abiotic environment. We have comparatively little knowledge about how polyploids interact with their surrounding biotic community (e.g., herbivores, pollinators, microbial symbionts; Forrester & Ashman, 2020; Forrester, Rebolleda-Gomez, Sachs, & Ashman, 2020; Segraves, 2017; Segraves & Anneberg, 2016), although species interactions are likely a major driver of polyploid establishment and persistence (Oswald & Nuismer, 2011). Evidence is accumulating that – compared to their diploid ancestors – polyploids can associate with a different subset of the local biotic community and even associate with novel species, such as mutualists (e.g., pollinators) or antagonists (e.g., herbivores) (Forrester & Ashman, 2020; Van de Peer, Ashman, Soltis, & Soltis, 2021; Streher et al. *In Prep*). These novel associations can, in turn, change the ‘macroscopic’ biotic community that plants associate with (Segraves, 2017).

A pivotal yet under-explored community-wide effect of polyploidy is the effect on the microbial community they host (Segraves, 2017; Segraves & Anneberg, 2016). Plant microbiomes consist of bacteria, fungi and viruses that live on (ectophytes) and within (endophytes) plant tissues (Cordovez, Dini-Andreote, Carrion, & Raaijmakers, 2019). While studies have demonstrated differential effects of polyploidy on individual root bacterial and fungal taxa (Anneberg & Segraves, 2019; Forrester & Ashman, 2018), characterization of the whole microbial communities of roots (rhizosphere) or leaves (phyllosphere) has lagged behind. Understanding the effect that whole genome duplication has on the microbiome is a crucial piece of missing information about their biotic interactions as well as our understanding of what drives the establishment of polyploid populations because the microbiome can influence plant performance (Bai et al., 2022; Gould et al., 2018; Parshuram, Harrison, Simonsen, Stinchcombe, & Frederickson; Tan, Kerstetter, & Turcotte, 2021; Yan, Levine, & Kandlikar, 2022).

Phenotypic changes due to polyploidy may cultivate novel assemblages of microbes (Doyle & Coate, 2019; Fox et al., 2020). For instance, a key phenotypic change is cell and organ enlargement (Doyle & Coate, 2019). Due to a greater surface area, the enlarged organs of polyploids may allow greater microbial colonization and, in turn, increased microbial diversity (Glassman, Lubetkin, Chung, & Bruns, 2017; Lyons et al., 2010; Peay, Garbelotto, & Bruns, 2010). Larger organs may also relax competition among potentially colonizing microbes (Ghoul & Mitri, 2016), allowing competitively inferior taxa to persist, thus increasing microbial diversity. Other key traits such stomatal size, chlorophyll content of cells, and secondary metabolite production could impact bacterial colonization or persistence. For example, neopolyploids of *A. thaliana* have larger stomates with a higher concentration of leaf chlorophyll (Yu, Haage, Streit, Gierl, & Ruiz, 2009), suggesting that neopolyploids could achieve greater metabolic potential than their diploid ancestors, thereby promoting the colonization and growth of microbes that associate with the leaf. Other studies have demonstrated that polyploidy can increase immune gene expression and secondary metabolite accumulation (Lavania et al., 2012; Song & Chen, 2015), suggesting that polyploidy may enhance the ability of plants to suppress or better tolerate pathogen colonization. Taken together, the many phenotypic novelties caused by neopolyploidy should also lead to novelty in the plant microbiome.

A handful of recent studies have shown that whole genome duplication has variable effects on plant microbial communities. Wipf & Coleman-Derr (2021) sequenced the bacterial rhizosphere of diploid, tetraploid, and hexaploid wheat species from a single field site and found that polyploid species hosted more microbial taxa (alpha diversity) and the composition of their root microbiome (beta diversity) was significantly differentiated from diploids. Similarly, synthetic tetraploids from two genotypes of *Arabidopsis thaliana* were exposed to a single inoculum source and found to host a greater abundance, but similar diversity, of bacterial taxa in the rhizosphere compared to their diploid ancestors (Ponsford et al., 2022). However, another experiment using seven synthetic tetraploid genotypes of *A. thaliana* found that polyploidy did not restructure the microbiome when exposed to a synthetic community 16 bacterial taxa but the polyploids were more tolerant to pathogenic attack than their diploid progenitors (Mehlferber et al., 2022). These latter two studies used synthetic “neopolyploids” (incipient polyploids), rather than established polyploids, and thus avoided the confounding effects of subsequent evolution following the whole genome duplication event, and yet they still observed variable responses to polyploidy, suggesting that the source microbial community might also determine the outcome. To determine the relative roles of neopolyploidy and inoculum source on the plant microbiome, we need studies that not only include multiple host genotypes, but also multiple inoculum sources. Indeed, we have yet to determine whether the ‘core microbiome’ – the set of microbial taxa that frequently colonize their host plants across a broad array of environmental contexts (Risely, 2020) – differs based on ploidy level.

The effect of neopolyploidy on the plant microbial community likely varies with genetic ancestry since polyploids often arise repeatedly from genetically different diploids (Soltis & Soltis, 1999; Soltis, Visger, Marchant, & Soltis, 2016), and these independent maternal origins can strongly influence the degree of phenotypic divergence from their diploid progenitors (Doyle & Coate, 2019). For example, Wei et al. (2020) used numerous genetically distinct diploid lineages of *Fragaria* to create synthetic neopolyploids and found that the effect of neopolyploidy on an array of functional traits was strongly dependent on diploid genetic ancestry. Similarly, Pacey et al. (2020) found substantial variation among 55 genetically diverse pairs of diploids and synthetic neopolyploids of *A. thaliana* in their expression of a set of leaf and physiological traits. Having increased genetic diversity among neopolyploid plants could impact the taxonomic composition of microbial communities since genotype is a strong predictor of microbiome assembly in many studies (Dastogeer, Tumpa, Sultana, Akter, & Chakraborty, 2020). For example, independent genetic lineages of neopolyploids can vary in their susceptibility to pathogens based on the allelic frequencies of resistance genes in their diploid ancestry (Oswald & Nuismer, 2007), which could lead to substantial variation among genetic lineages of polyploids in the abundance of pathogens that can colonize them (Van de Peer et al., 2021).

Thus, we expect that the effect of neopolyploidy on microbiome composition will vary across multiple genetic origins and by testing this assertion, we can more accurately measure the repeatable effect of polyploidy on the plant core microbiome. Variation in environmental setting (i.e., the abiotic and biotic parameters of a site) is likely to influence the effect of polyploidy on microbial communities (Eckert et al.; Li et al., 2023). In particular, the taxonomic diversity within inoculum can co-vary with the abiotic environment, such as with temperature and nutrient availabilities (Bakker, Chaparro, Manter, & Vivanco, 2015). Therefore, if polyploids preferentially associate with microbial taxa that are not present in certain environments, this could constrain the differences between diploid and neopolyploid microbiome communities. Moreover, if polyploid genotypes vary in their response to the environment (e.g., genetic variation among polyploid lineages in growth rate; Pacey et al., 2020), then environmental effects on the microbiome may be ploidy- or genetic lineage-dependent. In sum, the environmental context may mediate differences in microbial communities between polyploids and diploids, and possibly interact with their diploid ancestry (genotypic variation), yet no study has addressed the relative roles of whole genome duplication or genetic background on plant microbial community composition across a variety of ecological settings. Since polyploids may cultivate greater diversity in their microbiome communities than diploids, we expect that whole genome duplication will broaden the taxonomic core microbiome across these diverse ecological settings.

We tested how polyploidy affects plant bacterial communities using experimental inoculation of axenic diploids and synthetic neotetraploids of the aquatic plant “Greater Duckweed” *Spirodela polyrhiza* (L.) Schleid (Araceae). Duckweeds are quickly becoming a model system for studying the microbial ecology of plants (Baggs, Tiersma, Abramson, Michael, & Krasileva, 2022; Lam, Appenroth, Michael, Mori, & Fakhoorian, 2014; Tan et al., 2021), due in part to their simple morphology but also because their microbiomes are strongly filtered from their local aquatic habitat (Toyama et al., 2009). Additionally, their compact size and rapid generation time of 4 – 5 days (Acosta et al., 2021; Ziegler et al., 2015) makes duckweed an ideal system for testing how polyploidy affects microbial community assembly in laboratory settings.

We grew four genetically distinct pairs of diploids and their immediate neopolyploid descendants (Anneberg et al., 2023) individually in different water sources collected from 10 different ponds to answer the specific questions: 1) How does neopolyploidy affect the alpha and beta diversity of the duckweed bacterial community? 2) Does the effect of neopolyploidy on bacterial alpha and beta diversity depend more on diploid genetic ancestry or the ecological source of inoculum? 3) Compared to their diploid progenitors, is the core microbiome of neopolyploids taxonomically broadened or restricted? 4) If so, what are the key bacterial taxa that differentiate diploid versus neopolyploid core microbiome?

## Materials and Methods

### Study system

*Spirodela polyrhiza* is one of the largest duckweed species (Acosta et al., 2021), with vegetative fronds floating on the water surface and numerous simple roots growing into the water column. They have a cosmopolitan distribution (W. Wang, Kerstetter, & T.P., 2011), and naturally occur in freshwater habitats. We used colchicine-induced neo-autotetraploid and colchicine-exposed but unconverted diploids described in Anneberg et al., (2023) to test our questions. These neotetraploids (hereafter ‘neopolyploids’) came from four distinct diploid genotypes (hereafter ‘genetic lineages’) of *S. polyrhiza* collected in Western Pennsylvania, USA (Table S1). Previous work has shown that these neopolyploid lineages are larger than their diploid ancestors and vary in their growth rates, showing that neopolyploidy has led to phenotypic novelty in this system (Anneberg et al. 2023). To eliminate the microbial taxa that are sourced from the laboratory environment rather than natural settings, we generated axenic cultures of these duckweed lineages two weeks prior to the inoculation by following a modified protocol from Barks et al. (2018): Prior to surface-sterilization, we activated recalcitrant dormant microbes colonized on tissues by pre-culturing the duckweeds in half-strength Schenk-Hildebrandt media supplemented with 6.7 g/L sucrose, 0.067 g/L yeast extract, and 0.34 g/L tryptone powder for 24 hours (Schenk & Hildebrandt, 1972). We then submerged all pre-cultured tissues in 0.8% sodium hypochlorite solution in autoclaved deionized water (v/v) with gentle mixing using sterile forceps for six minutes before placing the sterilized plants in fresh quarter-strength Appenroth media (Appenroth, Teller, & Horn, 1996).

### Inoculum sourcing

We sampled inoculum as bulk water from ten ponds with large natural populations of *S. polyrhiza* spanning from upstate New York, USA to northern West Virginia, USA (Figure 1; Table S2). Therefore, our inoculum represents both biotic and abiotic attributes of these ponds. Between August 16^th^, 2021 to August 28^th^, 2021 we sampled three liters of raw pond water at each site from directly beneath a patch of wild duckweed near the shore by submerging two sterilized glass screwcap flasks fitted with 0.6 mm pore size mosquito netting over the opening to avoid collecting large solids. To profile the abiotic conditions of the inoculum source environments from where we collected water, we measured pH and took note of the site elevation and GPS coordinates (latitude and longitude). We additionally collected 500 mL of pond water from each pond source in the same way as described above and analyzed these samples for a set of abiotically relevant parameters that strongly influence microbial diversity (Dastogeer et al., 2020; Sadeghi, Chaganti, Shahraki, & Heath, 2021; Tang et al., 2020): total dissolved solids (TDS), phosphate, nitrate, iron, calcium carbonate, and sulfate at the Agriculture Analytical Services Lab in Pennsylvania State University. Last, as a control for contamination due to airborne microbes at sites or transportation of the water back to the lab, we opened a sterile 50 mL falcon tube filled with autoclaved millipore water and left this tube uncapped and exposed to the air for two minutes. We included one ‘field’ control for each of the ten sites. All samples collected from sites were placed into a cooler filled with ice and transported back to the University of Pittsburgh within 24 hours of collection.

**Figure 1:**
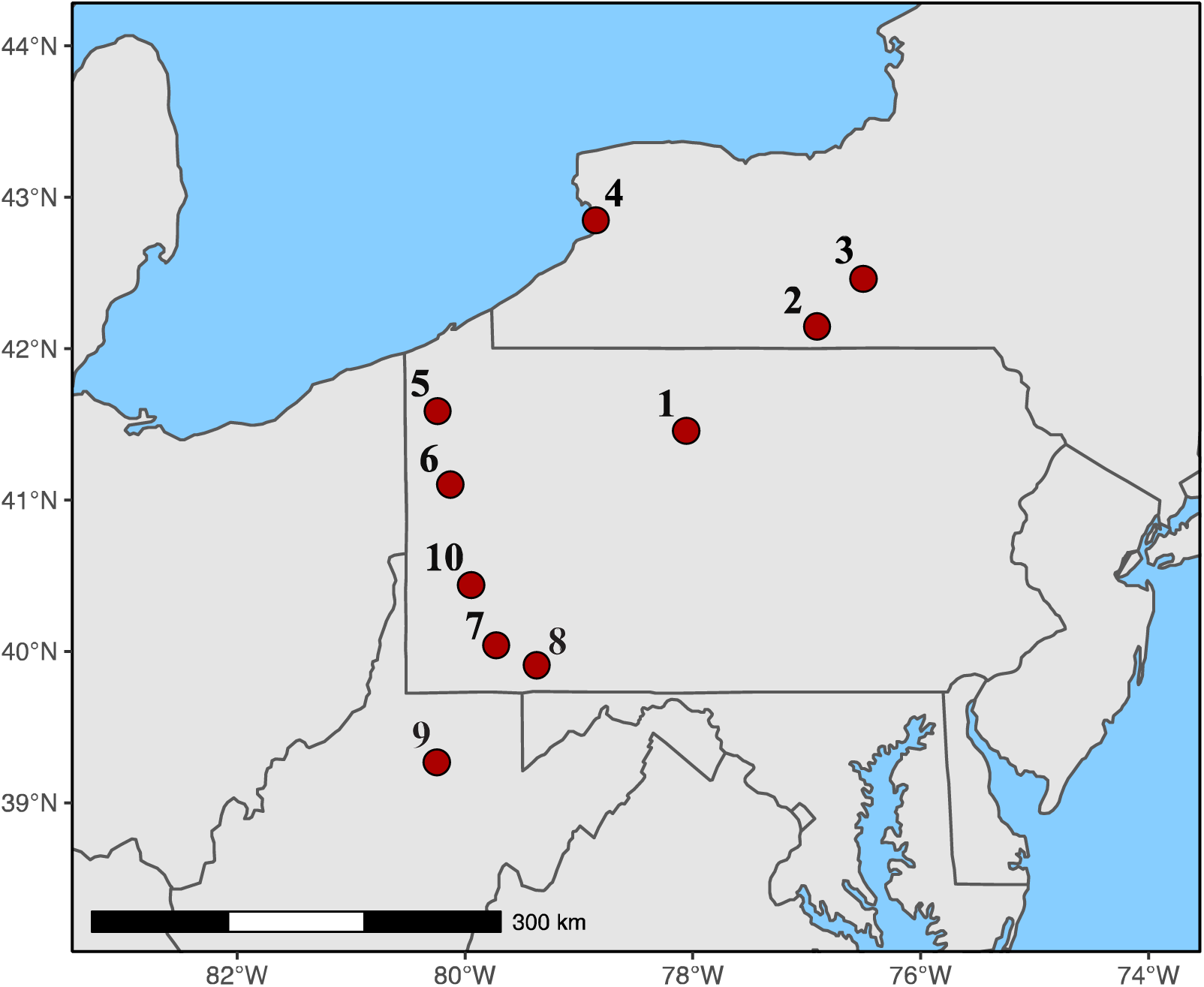
Map of the sources of inoculum in the northeastern United States. Source ponds correspond to the numbers supplementary table 1.

### Experimental Inoculation

To begin the inoculation experiment, for each pond source, we decanted 20 mL of that pond water into 48 sterile 25 mL glass test tubes (480 tubes total across all sites). Each tube then received three sterile *S. polyrhiza* fronds of a given ploidy level from a given genetic lineage. There were six replicates of each ploidy-genetic lineage-source combination. Due to the large geographic area of sampling, the experimental inoculation was conducted in three temporal blocks (Block One: 4 sources; Block Two: 3 sources; Block Three: 3 sources), all within a two-week period. To confirm that each lineage of surface-sterilized duckweed were axenic, we filled 48 tubes with 20 mL sterile half-strength Appenroth media for each temporal block (144 tubes total across three blocks). In total, we grew 624 *S. polyrhiza* samples distributed across two ploidy levels, four genetic lineages, ten inoculum water sources, plus six sterile ‘media’ controls for each genetic lineage by ploidy level and for each the three collection blocks (144 total). The media control served to evaluate bacterial taxa which are recalcitrant to our sterilization technique after generating axenic duckweed. To prevent environmental contamination but also allow air exchange, we capped each of the 25 mL glass tubes with an autoclaved gas-permeable sterile plastic cap. The tubes were placed in a completely randomized design in a growth chamber set to 16:8 L:D cycle with 25°C constant temperature and 50% relative humidity and incubated for two weeks, which was long enough for approximately three duckweed generations, until harvest.

We collected whole duckweeds for bacterial sequencing analysis by gently collecting plants with forceps via sterile technique and allowing them to drip dry on the sides of tubes. The drip-dried plant was then placed into a labeled 1.5 mL sterile centrifuge tube and stored in a - 80°C freezer until DNA could be extracted from whole plant tissue, fronds and roots included. Thus, the bacterial communities reported represent both endo- and ecotypes from both the rhizosphere and phyllosphere.

### Bacterial DNA sequencing and taxonomic identification

We randomly selected 4 – 5 of the six replicates of each ploidy-genetic lineage-inoculum source combination and extracted DNA from approximately 150 mg of tissue. DNA was extracted using Zymo Quick-DNA Fecal/Soil Microbe Miniprep (Zymo Research, Irving, CA, USA) and following manufacture’s protocol with the exception that the DNA elution buffer was increased in volume and time before centrifuging. In total, 352 duckweed samples and 19 media control samples were stored in a −20°C freezer until sequencing for the V5-V6 region of the bacterial 16s rRNA gene via the 799f and 1115r primer sets (Gilbert, Jansson, & Knight, 2014). One lane of paired-end sequencing on an Illumina MiSeq platform (Caporaso et al., 2012) was performed by Argonne National Lab (Illinois, USA).

A total of 15,692,711 reads were obtained and quality filtered and processed as follows. Using histograms of forward and reverse read quality scores and with the dada2 plugin (Callahan et al., 2016) in Qiime2 (Bolyen et al., 2019), we trimmed forward reads by 20 base pairs and reverse reads by 14 base pairs and truncated on length by 210 in the forward reads and 220 in the reverse reads. We next denoised the sequencing reads with default parameters of the dada2 plugin followed by generating a multiple sequence alignment tree with the fasttree Qiime2 function via mafft (Katoh & Standley, 2013). To assign taxonomy to each amplified sequence variant (ASV), we used the Qiime2 feature classifier plugin (Bokulich et al., 2018) with the Silva small subunit rRNA database release 138.1 (Quast et al., 2013). To merge the ASV counts table, multiple alignment tree, taxonomy table, and metadata files, we imported these files into R (R Core Team, 2021) with the phyloseq package (McMurdie & Holmes, 2013). Since one of the kit controls did not amplify any sequences, the dada2 pipeline removed that sample from the dataset, and the resulting phyloseq object was comprised of 10,365 ASVs across 370 samples. We filtered out mitochondrial or chloroplast DNA and this removed 1,768 ASVs, leaving 8597 bacterial ASVs in the dataset. We further filtered the phyloseq data by removing any ASVs that were detected in either our media controls or field water controls from each collection site. This removed another 189 ASVs, leaving 8408 ASVs from 320 samples. The relative taxonomic abundance of these 189 ASVs that were detected in our sterile duckweed (media controls) are presented in Fig S1, and a full taxonomy table of these taxa is freely available as Appendix S1. Our final filtering step involved applying a loss function to the ASV table to remove spurious taxa that were rarely detected and could be the product of sequencing error or misclassification. Rather than applying an arbitrary abundance threshold cutoff, we used the PERFect package (Smirnova et al., 2019) in R to permutationally filter out rare ASVs that do not have a detectable effect on the entire ASV covariation matrix (Smirnova et al., 2019). By passing our phyloseq object through PERFect filtering, we removed 5,122 ASVs that were in insignificantly low read abundance in the dataset, resulting in a final filtered phyloseq object comprised of 3,286 ASVs across 320 experimentally inoculated duckweed samples. In this final phyloseq dataset, we normalized the reads per sample to the overall median (Wei & Ashman, 2018).

### Statistical Analysis

We quantified the effect of ploidy level, genetic lineage, and inoculum source on the alpha diversity of the duckweed bacterial community. To calculate bacterial alpha diversity, we used the phyloseq function “estimate_richness” to determine taxonomic (ASV) richness and Shannon diversity. We calculated Faith’s phylogenetic diversity with the function “pd”. Because we started the inoculation experiment in three separate blocks spread over two weeks, we tested for a temporal block effect in our statistical analyses. No block effect was observed in diversity analyses and is therefore not reported further. All statistical analyses were performed with R software (R Core Team, 2021), using the base stats functions as well as the packages ampvis2 (Andersen, Kirkegaard, Karst, & Albertsen, 2018), edgeR (Robinson, McCarthy, & Smyth, 2010), lsmeans (Lenth, 2016), picante (Kembel et al., 2010), and vegan (Dixon, 2003). Each alpha diversity metric was analyzed with a linear model, defining ploidy level, genetic lineage, and inoculation source as main effects, along with their interaction terms.

To estimate beta diversity of duckweed bacterial communities, we analyzed Bray-Curtis dissimilarities. We calculated Bray-Curtis dissimilarities with the distance function in the phyloseq package and then used the Adonis2 function from the vegan package to fit a permutational analysis of variance (PERMANOVA) model to the data. In the PERMANOVA, we defined the model as: Bray-Curtis dissimilarities explained by ploidy level, genetic lineage, inoculation source, and their interaction terms. We then tested for homogeneity of variance among the groups in the PERMANOVA model by measuring beta dispersion for each factor via the betadisper function in the vegan package. We further assessed whether bacterial community compositions of neopolyploids are more likely to resemble each other instead of diploids by conducting a nearest neighbor analysis of the Bray-Curtis dissimilarities of samples. We did so by constructing a minimum spanning tree (Friedman & Rafsky, 1979) which aligns samples based on their Bray-Curtis dissimilarities. To test whether nearest neighbors of samples belong to the same groups or not, we conducted a permutational graph-based test by using the graph_perm_test function from the phyloseqGraphTest package (Fukuyama, 2020) in R with 10,000 permutations.

To infer which bacterial ASVs contribute to the differentiation of the diploid and neopolyploid communities, we built a random forest model (Breiman, 2001) with the randomForest function and specifying 2000 trees. Using this model, we classified diploid versus neopolyploid microbiomes given the differences in their ASV communities. We recorded the out of bag error rate of the overall model and the specific class errors between diploids and neopolyploids respectively with the confusion matrix. From the random forest model and using the “importance” function in R, we derived a list of the 30 most important ASVs that discriminates diploid versus neopolyploid microbiomes based on the mean decrease in the Gini impurity index (Nembrini, Konig, & Wright, 2018) for each ASV. We further tested for key taxa differentiating neopolyploids from their diploid progenitors by quantifying the log_10_-fold change of the most differentially abundant ASVs between diploids and neopolyploids. To do so, we first selected only the ASVs with variance in their counts below a threshold of 1*10^-6^ and used the phyloseq_to_edgeR function from McMurdie & Holmes (2013) to carry out an exact test of ASV abundance between diploids and neopolyploids using the exacttest function in the edgeR package (Robinson et al., 2010). We then adjusted p-values from multiple testing following Benjamini & Hochberg (1995) while implementing the topTags function in the edgeR package.

Last, we assessed the effect of polyploidy on the composition of the core microbiome of *S. polyrhiza* via the “amp_venn” function. We used a minimum 0.1 % relative abundance as a cutoff for being included in the core. After accounting for the relative abundance cutoff, the core bacterial microbiome of diploid and neopolyploid duckweed was characterized with a frequency cutoff of 50%, meaning that any “core” ASV must be colonized in at least 50% of all samples within any one of the three possible ploidy groupings (2x exclusive, 2x and 4x combined, and 4x exclusive core microbiome). However, we also report on a second, more conservative relative frequency cutoff of ≥ 75% to evaluate how restricted the core microbiome estimation can be to relative frequencies within groups.

## Results

### Polyploidy increases the alpha diversity of duckweed bacterial microbiome

Neopolyploids had significantly greater bacterial alpha diversity than their diploid progenitors across all three diversity measures (Table 1): Polyploidy increased taxonomic diversity by 4-11% (Shannon diversity and species richness, respectively) and phylogenetic diversity by 8% (Faith’s PD) (Fig. 2). Despite the alpha diversity of the bacterial microbiome of duckweed significantly depending on the inoculum source water, it did not interact with ploidy level (Table 1).

**Table 1:** Results of ANOVAs of the effects of ploidy, genetic lineage, inoculation source and their interactions on taxonomic diversity (Species richness and Shannon diversity) and phylogenetic diversity (Faiths’ PD) of bacterial communities of duckweed. Significant factors in are in bold. There were 240 residual degrees of freedom across all three alpha diversity models.

| Factor | Df | Shannon diversity |  | Species richness |  | Faith's PD |  |
| --- | --- | --- | --- | --- | --- | --- | --- |
|  |  | F value | P value | F value | P value | F value | P value |
| Ploidy | 1 | 6.97 | **0.009** | 6.22 | **0.013** | 5.79 | **0.017** |
| Lineage | 3 | 2.03 | 0.111 | 1.38 | 0.249 | 1.78 | 0.152 |
| Inoculum Source | 9 | 2.59 | **0.007** | 1.71 | 0.088 | 3.95 | **<0.001** |
| Ploidy: Lineage | 3 | 0.24 | 0.866 | 0.16 | 0.924 | 0.14 | 0.935 |
| Ploidy: Inoculum Source | 9 | 0.86 | 0.562 | 0.30 | 0.975 | 0.44 | 0.910 |
| Lineage: Inoculum Source | 27 | 0.66 | 0.903 | 0.72 | 0.840 | 0.73 | 0.830 |
| Ploidy: Lineage: Inoculum Source | 27 | 0.77 | 0.793 | 0.68 | 0.880 | 0.81 | 0.740 |

**Figure 2:**
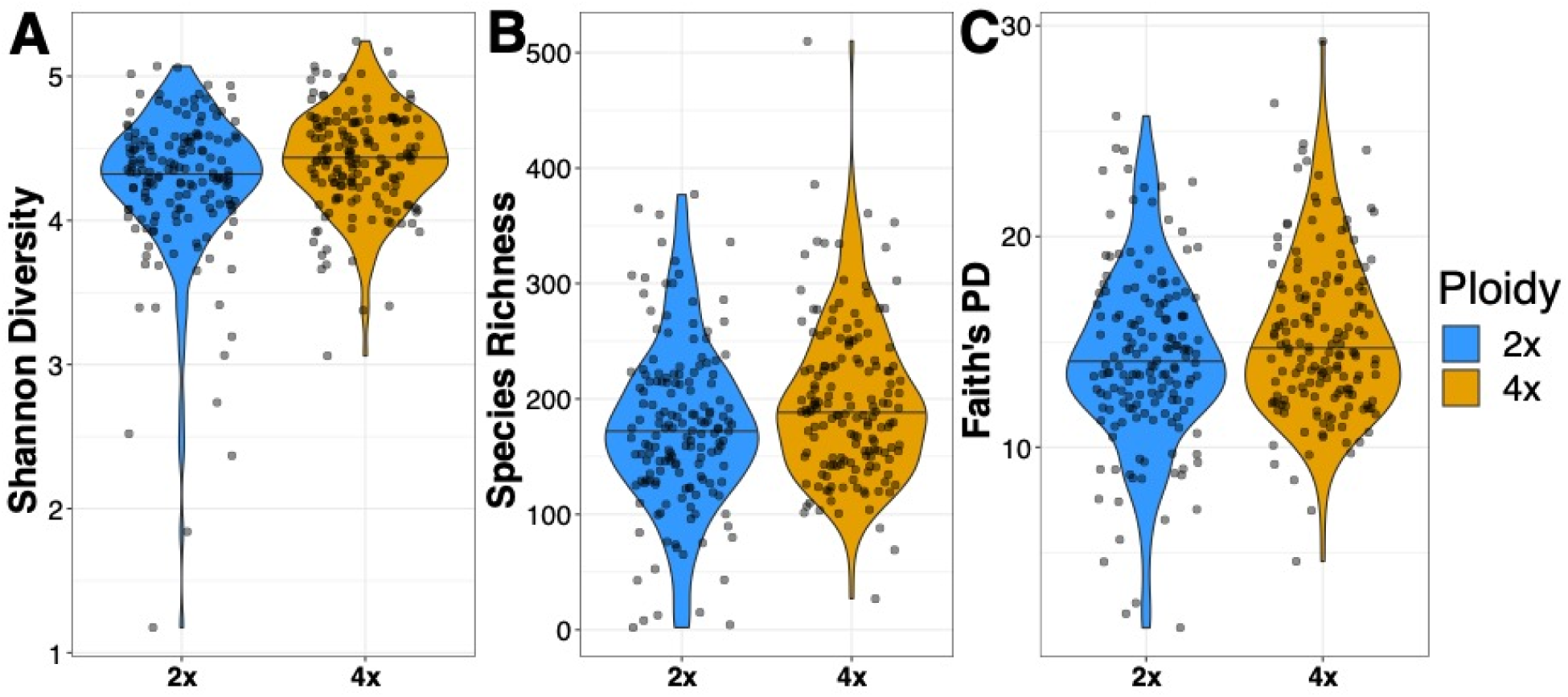
Violin plots showing the median and distribution of alpha diversity of bacterial communities for diploids (2x) and neopolyploids (4x). Neopolyploids had greater alpha diversity than their diploid progenitors for A) Shannon diversity, B) Species richness, and C) Faith’s phylogenetic distance. The median is denoted by a lateral black line in each violin plot and independent samples are represented by each black point plotted over the violin plots.

### The effect of polyploidy on duckweed microbiome composition is environment and genetic background dependent

The effect of polyploidy on beta diversity of duckweed bacterial communities was influenced by genetic lineage and inoculum source (F_27,240_ = 1.14, P = 0.029; Table 2; Fig. 3). This three-way interaction between ploidy, genetic lineage, and inoculum source explained approximately 4 % of the total variation in Bray-Curtis dissimilarities among samples (Table 2). Beyond this complex three-way interaction, there was a polyploidy by environment interaction that explained approximately 2% of the variation in the model as well as an interaction between polyploidy and genetic lineage explaining an additional 1% of variation (Table 2).

**Figure 3:**
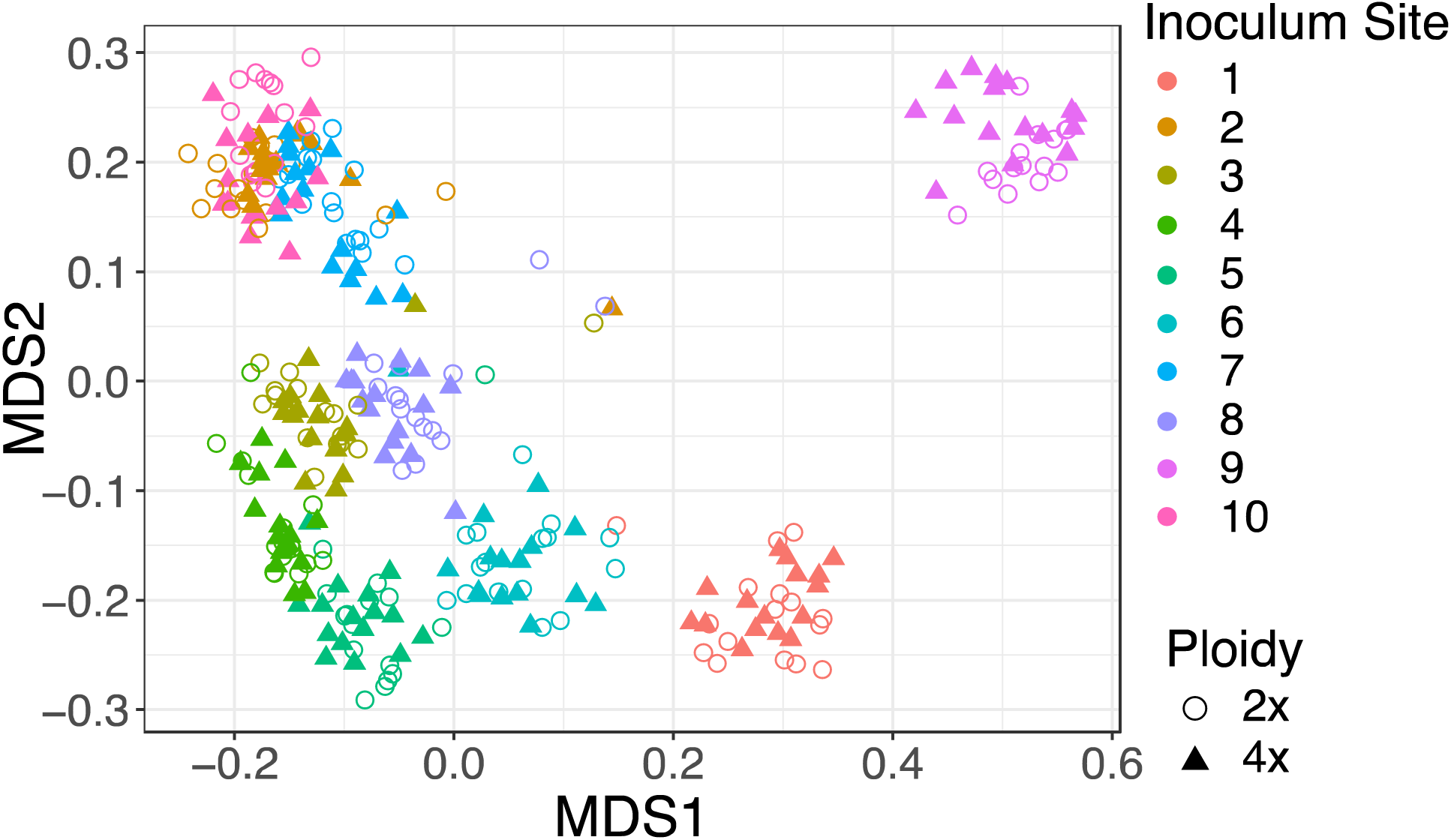
PCoA plot of bacterial Bray-Curtis dissimilarities colonized on diploid (2x) and neopolyploid (4x) duckweeds. Each point represents an independent duckweed plant and is colored by the site of the inoculum source (number represent the locations in Figure 1) and shapes represent ploidy level.

**Table 2:**
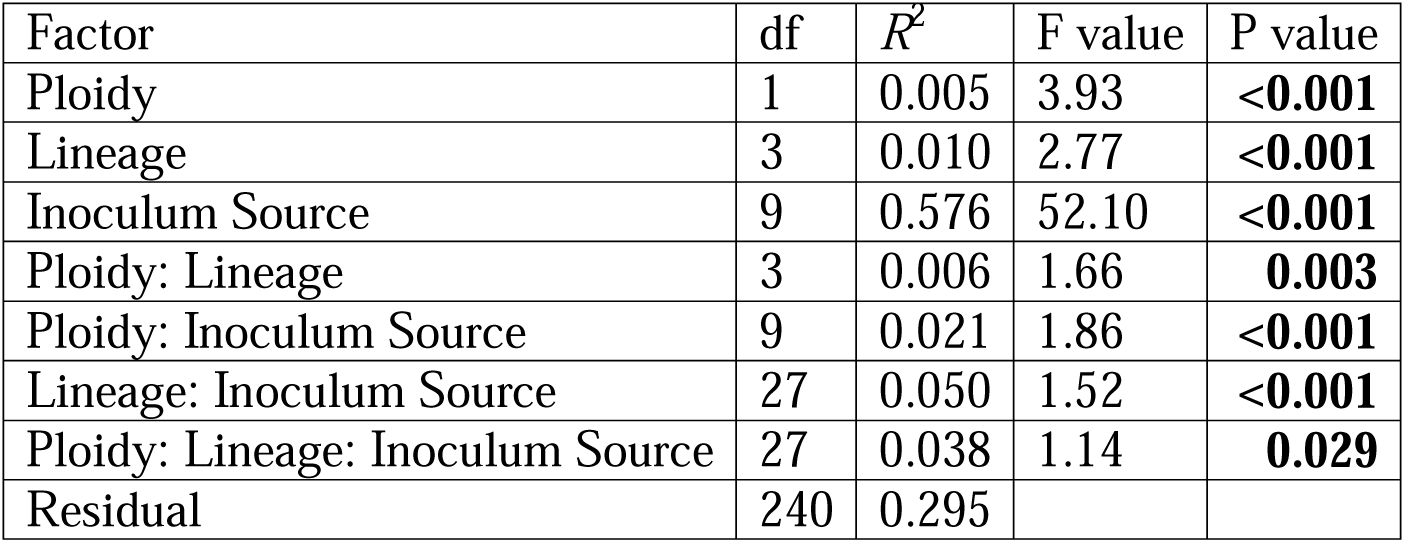
Results from PERMANOVA to evaluate the effect of ploidy, genetic lineage, inoculation source and their interactions on beta diversity of duckweed bacterial communities (Bray-Curtis dissimilarities). Significant effects in the model have a bolded P value.

The minimum spanning tree showed that neopolyploid bacterial communities resemble each other significantly more than they resemble the bacterial communities of their diploid ancestors (P < 0.0001; Fig. 4A). Similarly, samples inoculated with the same pond water resembled each other significantly more than a random neighbor in Bray-Curtis dissimilarity (P < 0.0001 Fig. 4B).

**Figure 4:**
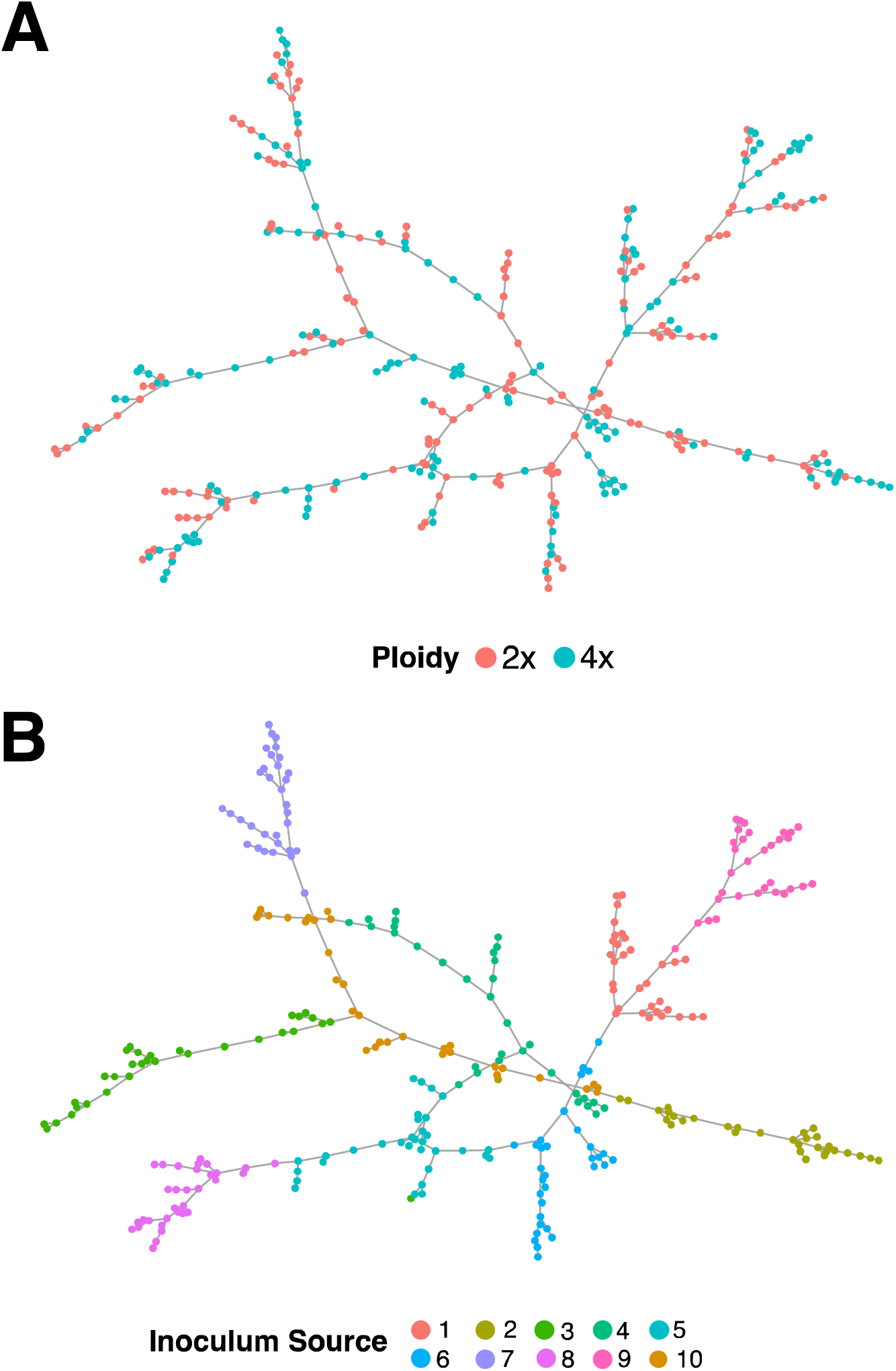
Minimum spanning trees that align samples of bacterial communities according to their nearest neighbor in Bray-Curtis dissimilarity, showing when they are colored by A) the ploidy level of the *S. polyrhiza* host plant (diploid-2x; nepolyploid-4x), and B) the inoculum source (number represent the locations in Figure 1) that duckweeds were inoculated with at the beginning of the experiment.

### Key taxa differentiate diploid and neopolyploid duckweed microbiomes

Several bacterial taxa uniquely colonize neopolyploids versus diploids. The random forest model that classified the ploidy level of samples based on the Bray-Curtis dissimilarities among samples had an out-of-bag error rate of 27 %, meaning that the model could correctly identify whether a sample bacterial community belongs to either a diploid or neopolyploid 73 % of the time. This random forest model was not biased in the ability to correctly identify a diploid bacterial community from a neopolyploid community, as indicated by the accompanying confusion matrix showing that the class-wise error rates were approximately equal for both diploid and neopolyploid bacterial communities (Table S3). The 30 indicator ASVs that discriminate diploid and neopolyploid *S. polyrhiza* were primarily members of the families Rhizobiaceae (n = 7) and Comamonadaceae (n = 6) in the Alphaproteobacteria and Gammaproteobacteria, respectively (Fig. 5; Table 3).

**Figure 5:**
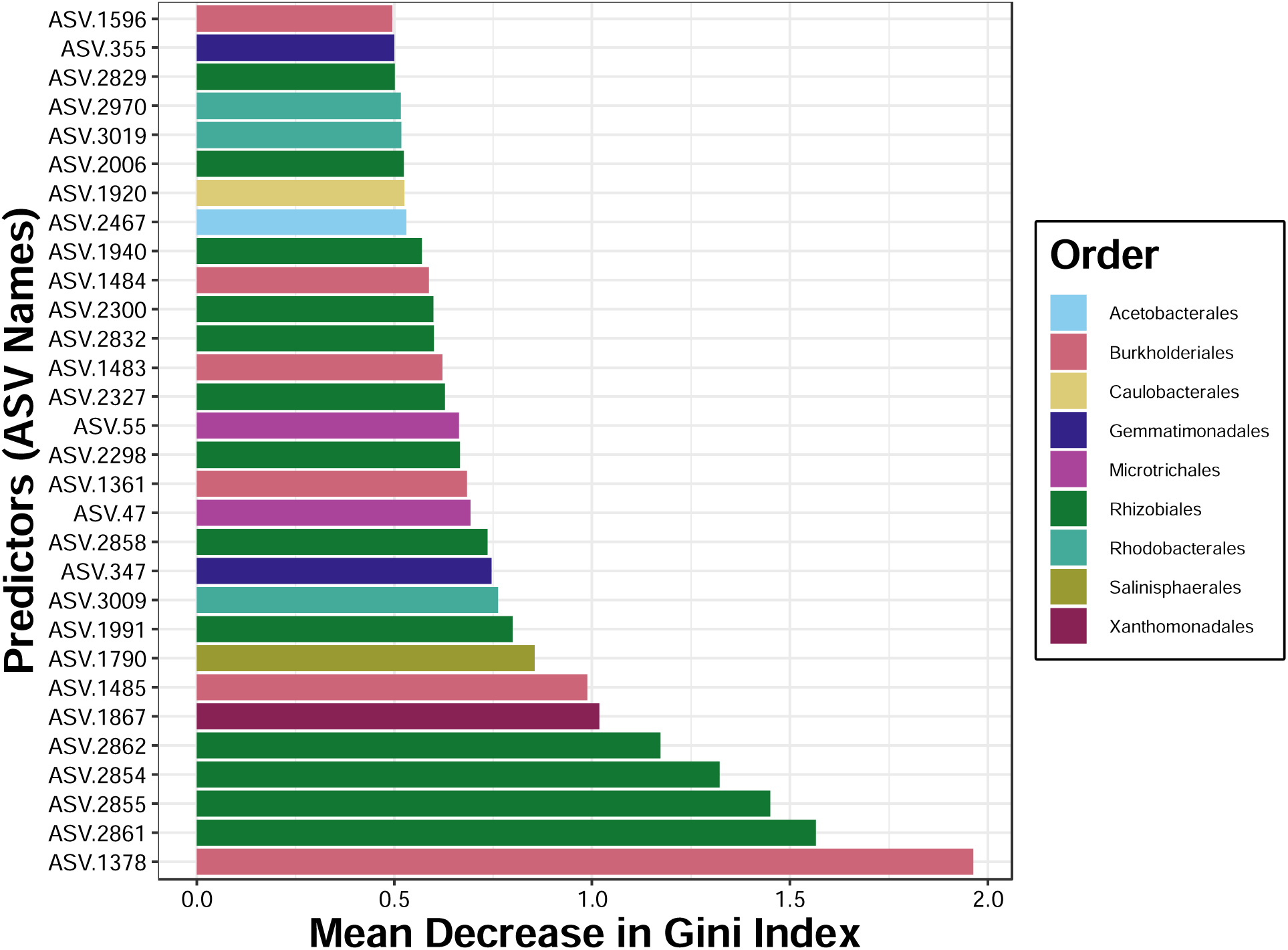
Gini index plot which shows the most important bacterial ASVs delineating the bacterial communities of diploids from neopolyploids in the random forest model. The greater the mean decrease in the Gini index, the more important that ASV is in discriminating diploids from neopolyploids. Individual bars are colored by their taxonomic order and the ASV names correspond to the ASV number in the taxonomy table in Table 3.

**Table 3:** The 30 most important bacterial genera (or family) that discriminate microbiome composition of diploid and neopolyploid duckweeds based on a random forest model of ASVs. ASV designation corresponds to the ASV predictors Gini importance index values in Fig S1. NA under the genus column represents ASVs that were not identifiable at the genus level.

| ASV Name | Phylum | Class | Order | Family | Genus |
| --- | --- | --- | --- | --- | --- |
| ASV.47 | Actinobacteriota | Acidimicrobiia | Microtrichales | Ilumatobacteraceae | NA |
| ASV.55 | Actinobacteriota | Acidimicrobiia | Microtrichales | Ilumatobacteraceae | Ilumatobacteraceae |
| ASV.347 | Gemmatimonadota | Gemmatimonadetes | Gemmatimonadales | Gemmatimonadaceae | Gemmatimonas |
| ASV.355 | Gemmatimonadota | Gemmatimonadetes | Gemmatimonadales | Gemmatimonadaceae | Gemmatimonas |
| ASV.1361 | Proteobacteria | Gammaproteobacteria | Burkholderiales | Comamonadaceae | NA |
| ASV.1378 | Proteobacteria | Gammaproteobacteria | Burkholderiales | Comamonadaceae | NA |
| ASV.1483 | Proteobacteria | Gammaproteobacteria | Burkholderiales | Comamonadaceae | NA |
| ASV.1484 | Proteobacteria | Gammaproteobacteria | Burkholderiales | Comamonadaceae | NA |
| ASV.1485 | Proteobacteria | Gammaproteobacteria | Burkholderiales | Comamonadaceae | NA |
| ASV.1596 | Proteobacteria | Gammaproteobacteria | Burkholderiales | Comamonadaceae | Rhizobacter |
| ASV.1790 | Proteobacteria | Gammaproteobacteria | Salinisphaerales | Solimonadaceae | Nevskia |
| ASV.1867 | Proteobacteria | Gammaproteobacteria | Xanthomonadales | Xanthomonadaceae | Silanimonas |
| ASV.1920 | Proteobacteria | Alphaproteobacteria | Caulobacterales | Hyphomonadaceae | Hirschia |
| ASV.1940 | Proteobacteria | Alphaproteobacteria | Rhizobiales | Devosiaceae | Devosia |
| ASV.1991 | Proteobacteria | Alphaproteobacteria | Rhizobiales | Pleomorphomonadaceae | Chthonobacter |
| ASV.2006 | Proteobacteria | Alphaproteobacteria | Rhizobiales | Pleomorphomonadaceae | Chthonobacter |
| ASV.2298 | Proteobacteria | Alphaproteobacteria | Rhizobiales | Hyphomicrobiaceae | Pedomicrobium |
| ASV.2300 | Proteobacteria | Alphaproteobacteria | Rhizobiales | Hyphomicrobiaceae | Hyphomicrobium |
| ASV.2327 | Proteobacteria | Alphaproteobacteria | Rhizobiales | Hyphomicrobiaceae | Hyphomicrobium |
| ASV.2467 | Proteobacteria | Alphaproteobacteria | Acetobacterales | Acetobacteraceae | Roseomonas |
| ASV.2829 | Proteobacteria | Alphaproteobacteria | Rhizobiales | Rhizobiales | uncultured |
| ASV.2832 | Proteobacteria | Alphaproteobacteria | Rhizobiales | Rhizobiales | uncultured |
| ASV.2854 | Proteobacteria | Alphaproteobacteria | Rhizobiales | Rhizobiaceae | Allorhizobium-Neorhizobium |
| ASV.2855 | Proteobacteria | Alphaproteobacteria | Rhizobiales | Rhizobiaceae | NA |
| ASV.2858 | Proteobacteria | Alphaproteobacteria | Rhizobiales | Rhizobiaceae | NA |
| ASV.2861 | Proteobacteria | Alphaproteobacteria | Rhizobiales | Rhizobiaceae | Allorhizobium-Neorhizobium |
| ASV.2862 | Proteobacteria | Alphaproteobacteria | Rhizobiales | Rhizobiaceae | NA |
| ASV.2970 | Proteobacteria | Alphaproteobacteria | Rhodobacterales | Rhodobacteraceae | NA |
| ASV.3009 | Proteobacteria | Alphaproteobacteria | Rhodobacterales | Rhodobacteraceae | NA |
| ASV.3019 | Proteobacteria | Alphaproteobacteria | Rhodobacterales | Rhodobacteraceae | NA |

For the bacterial taxa shared by diploid and neopolyploid duckweed, we found 90 ASVs that were significantly differentially abundant between the two ploidy levels (Fig. 6). Of these 90 ASVs, 63 of them were members of the Proteobacteria. The most common taxonomic families of the differentially abundant ASVs were 23 members of the Comamonadaceae and 11 members of the Rhizobiaceae. Neopolyploids tended to have greater abundance of these ASVs than diploids (Fig. 6).

**Figure 6:**
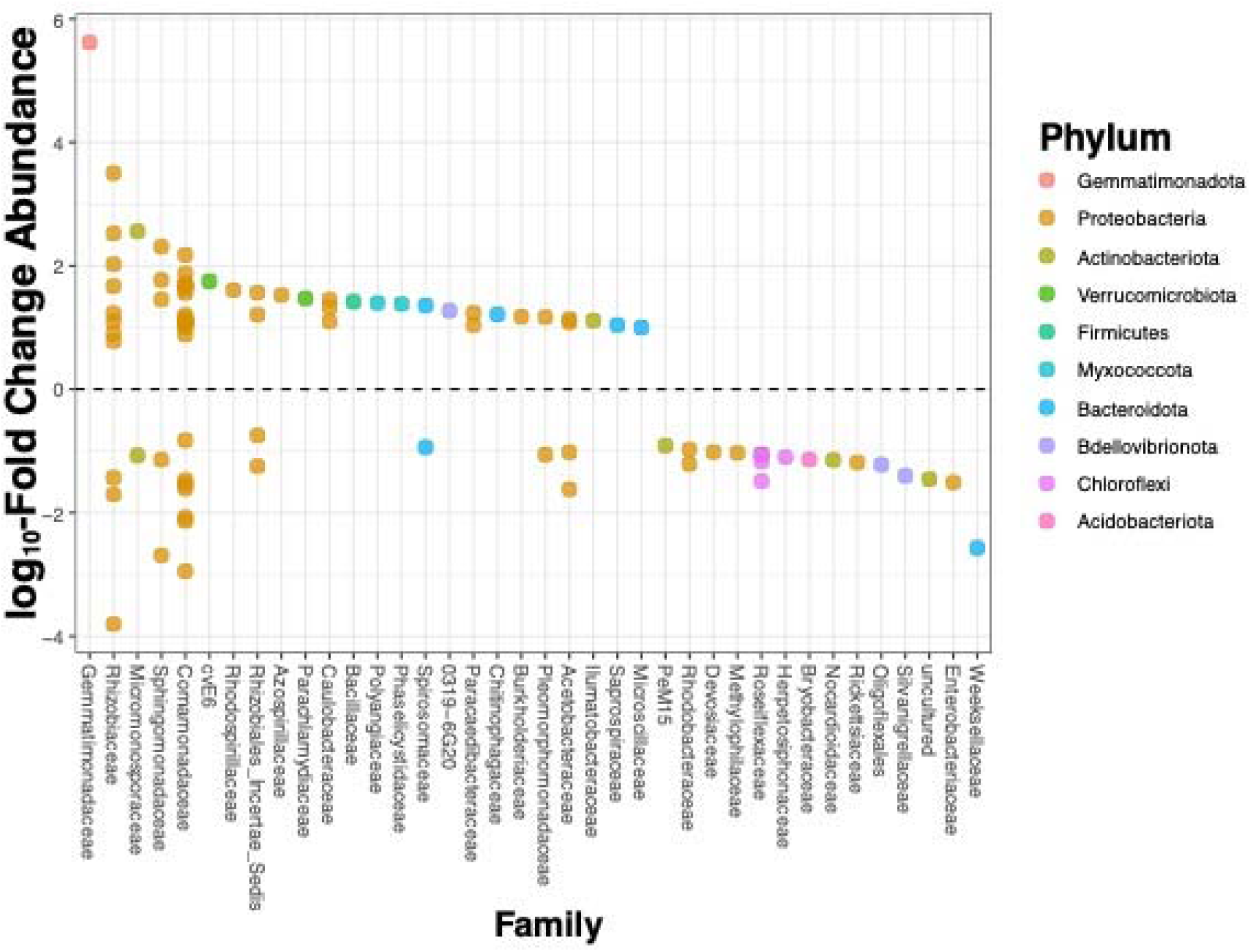
Log_10_-fold change in the most differentially abundant bacterial ASVs (grouped by family on the x-axis) between diploid and neopolyploid *S. polyrhiza*. Any ASV denoted with a colored dot above zero means that was found more often on neopolyploids than diploids, and the reciprocal applies to any ASV plotted below zero.

### Polyploidy broadens the core microbiome

When the core microbiome was characterized by a 50% frequency cutoff, there were 55 bacterial ASVs shared between diploids and neopolyploids, 15 ASVs uniquely hosted by neopolyploids, and none were unique to diploids (Fig 7A, Appendix S2). Increasing the stringency of the frequency cutoff defining the core to 75% yielded 15 shared core ASVs between diploids and neopolyploids, and 5 ASVs that were unique to neopolyploids (Fig. 7B). At this level of stringency, the majority of the shared bacteria were members of the Proteobacteria belonging to either the Burkholderiales or Rhizobiales order (Appendix S2). Although we were unable to resolve the taxonomy of the five ASVs belonging to the neopolyploid-exclusive core bacterial more specific than the family level, one was a member of the Actinobacteriota phylum belonging to the Illumatobacteriacea family, and the other four were members of the Proteobacteria belonging to the families: Comamonadaceae, Rhizobiales (*Incertae sedis*) Solimonadaceae, Xanthomonadaceae (Appendix S2).

**Figure 7.**
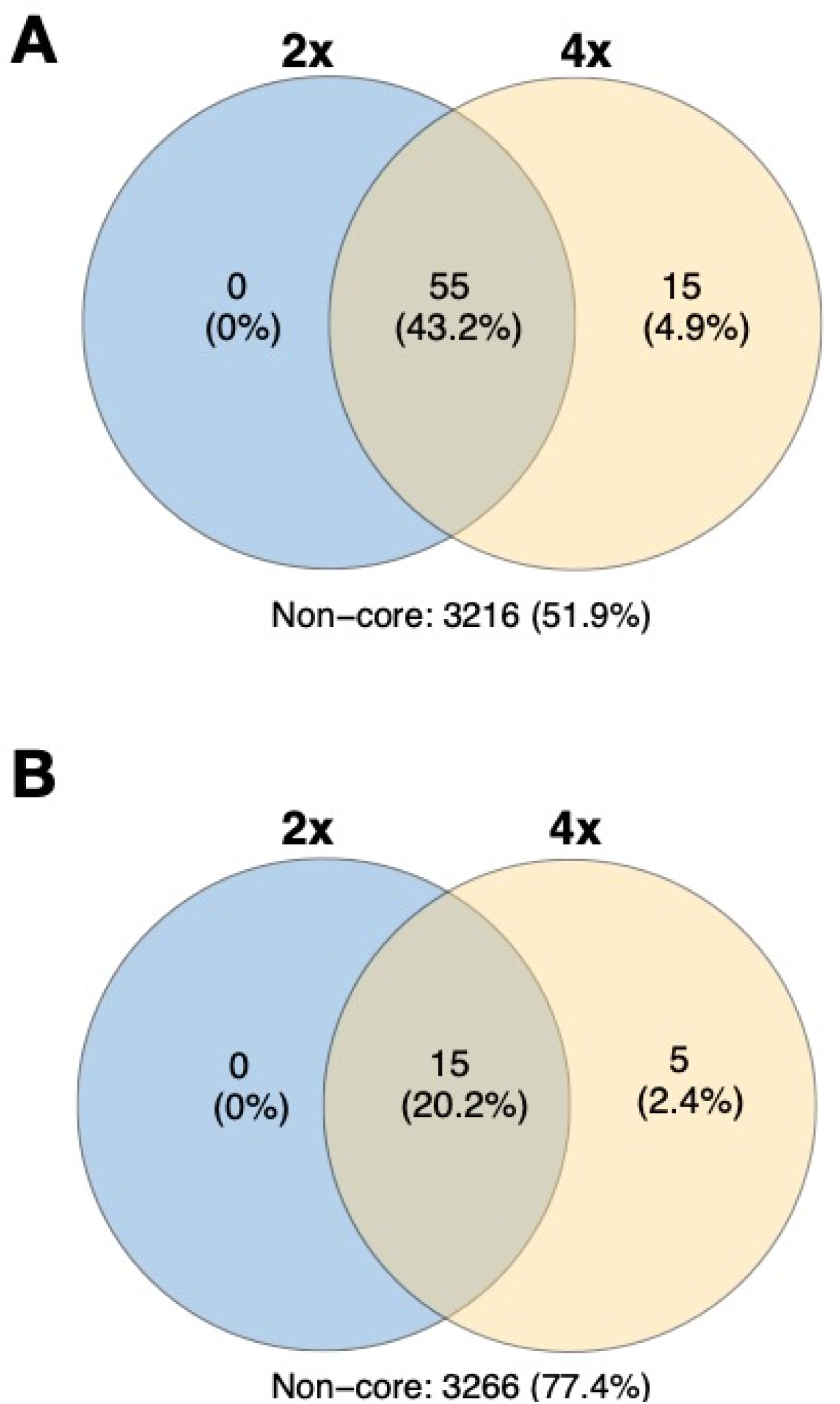
Venn diagrams of the number of bacterial ASVs either shared between diploids (2x) and neopolyploid (4x) duckweeds or uniquely hosted by them in the core microbiome. Number of core bacterial ASVs when we used a frequency cutoff across communities of diploids, neopolyploids, or both of either 50% (A) or 75% (B) is used.

## Discussion

Our culture-independent test of how neopolyploidy affects the duckweed microbiome across a broad set of naturally sourced inoculum revealed that ploidy level, genetic lineage, inoculum source, and their interactions play major roles in the diversity and composition of the plant microbiome (Table 2). Although diploid and neopolyploid microbiomes were similar when inoculated with some pond water sources, our minimum spanning tree analysis showed that the nearest neighbors in bacterial community composition were of the same ploidy level more often than not (Fig. 4). This suggests that the signature of polyploidy on bacterial community composition is observable across diverse natural aqueous environments which encompasses variability in both abiotic and biotic diversity. Furthermore, we found that polyploidy increases the taxonomic and phylogenetic diversity of the bacterial community as well as broadening the taxonomic core microbiome of *S. polyrhiza*. The neopolyploid core microbiome is comprised of novel taxa that rarely colonize their diploid progenitors while retaining the taxonomic core members of their diploid progenitors. Thus, our results show that polyploidy can immediately broaden the biotic niche of plants, and importantly that this effect is dependent on both genetic ancestry as well as the ecological setting that neopolyploids arise in.

Our manipulative experiment found that the effect polyploidy on the plant bacterial microbiome is generalizable over multiple genetic origins and across a variety of pond water sources. While our work also corroborates similar previous work (Ponsford et al., 2022; Wipf & Coleman-Derr, 2021) in that neopolyploidy increased bacterial alpha diversity (Fig. 2) and differentiated the plant bacterial community structure (Fig. 3), we additionally found that this effect varied among the sources of inoculum as well as the multiple genetic lineages of neopolyploids that we used (Table 2). Since the total diversity within the bacterial microbiome increased with neopolyploidy and this effect did not interact with inoculum source or genetic background (Table 1), this suggests that polyploidy has a universal effect on the alpha diversity of the plant bacterial microbiome. In contrast, both the source of inoculum water from as well as genetic lineage strongly interacted with polyploidy in structuring the beta diversity of the plant microbiome (Table 2), revealing that genetic ancestry and the ecological setting in which polyploids arise can have a deterministic effect of whether polyploidy differentiates bacterial community compositions from their diploid progenitors. The fact that some pond water sources did not lead to compositional differences in neopolyploid microbiomes from their diploid progenitors implies that these habitats could have an increased risk of neopolyploid extinction due to stronger niche overlap with diploids (Fowler & Levin, 2016; Rodriguez, 1996).

Conversely, the pond environments in which neopolyploids had strongly differentiated bacterial microbiomes shows a broadened biotic niche which could improve the odds of neopolyploid establishment through relaxing the competitive forces with their diploid ancestors. Our analysis of the taxonomic core microbiome – the bacterial taxa that frequently colonize host plants across broad ecological contexts (Risely, 2020) – showed that neopolyploidy broadened the core microbiome of *S. polyrhiza* (Fig. 7). Following previous studies that have used a 50 % frequency cut-off in determining the core versus non-core microbiome taxa (Ainsworth et al., 2015; Vidal-Verdu, Gomez-Martinez, Latorre-Perez, Pereto, & Porcar, 2022), we found that neopolyploid core microbiome was comprised of an additional 15 taxa on top of the 55 taxa they share with the diploid core microbiome (Fig. 7). This result reveals that neopolyploidy leads to increased generalism in the symbiosis between *S. polyrhiza* and bacteria. We can draw analogies of our findings to previous work that characterized how pollinator communities are influenced by polyploidy, such as in *Chamerion angustifolium,* in which diploids and polyploids strongly overlap in the pollinator communities that they host, but the polyploids are visited by two additional pollinator species that do not visit diploids (Kennedy, Sabara, Haydon, & Husband, 2006). Taken together, our results show that neopolyploidy leads to immediate novelty in the biotic niche of plants.

Neopolyploids cultivated a greater abundance of key bacterial taxa in their microbiome compared to diploids. Similar to previous work that has characterized the bacterial microbiome of duckweeds, we found that members of the Proteobacteria were dominant across all duckweed samples (Acosta et al., 2020; Bunyoo et al., 2022; Iwano et al., 2020; Iwashita et al., 2020). However, polyploidy not only increased the relative abundance of these key bacterial taxa (Fig. 6), but it also increased the phylogenetic diversity of bacteria that colonize duckweed (Fig. 1), again demonstrating that polyploidy broadens the biotic niche. Although our 16s sequencing data could not resolve taxonomy below either the family or genus level, we found that polyploids host particularly more *Rhizobiaceae* and *Comamonadaceae* than their diploid progenitors (Table 3; Fig. 6). The *Rhizobiaceae* are noteworthy for their diverse function in influencing plant growth, as some taxa within this family are growth-promoting for provisioning nitrogen to host plants while others can hinder plant growth though enhancing pathogen attack (Carareto Alves, de Souza, Varani, & Lemos, 2014). Two key previous studies that have characterized the microbiome of duckweed have specifically found that duckweed have an increased representation of nitrogen-fixing bacterial members of the genus *Rhizobium* within the *Rhizobiaceae* compared to the surrounding environment (Acosta et al., 2020; Zhao et al., 2015). While our data set did not have the taxonomic precision to test for differences between diploids and neopolyploids in the colonization of nitrogen-fixing bacteria, the significantly greater colonization of members of the Rhizobiaceae on polyploids warrants future metagenomic studies that could achieve that level of precision.

By using multiple genetic lineages of axenic diploids and neopolyploids inoculated with a variety of pond water sources, we revealed that whole genome duplication can immediately expand the plant biotic niche. In particular, polyploidy not only increased the diversity and restructured the composition of the bacterial microbiome, but it also caused many of the dominant taxa in the duckweed microbiome to increase in relative abundance, suggesting that neopolyploidy enhances the quality of the host plant for these taxa. Future work that seeks to understand the factors which promote polyploid establishment should consider novelty in species interactions as a potential mechanism of polyploid persistence.

## Acknowledgements

We thank Elizabeth O’Neill for the generation and maintenance of the synthetic neopolyploid lineages, Jason Simmons for assistance in field collection and media preparation, and Nevin Cullen for feedback on experimental design and data analysis. This project was supported by the Dietrich School of Arts & Sciences and grants from the National Science Foundation (#2109452 to T.J.A., #2027604 & 1912180 to T-L.A., and #1935410 to M.M.T.).

## Data Accessibility

All 16s rRNA gene sequences generated from this work are deposited into the NCBI SRA database under the accession ID PRJNA946155. We also deposited all underlying data in the production of this article to be made freely available from the Zenodo data repository (DOI: 10.5281/zenodo.7750374).

## Benefit-Sharing

The benefits generated from this work accrue from our freely available data on public data repositories described above.

## Author Contributions

TJA, TLA, and MMT conceptualized and designed the experiment. TJA collected field and laboratory samples, carried out the experiment, analyzed the data, and wrote the first draft of the manuscript. TJA, MMT, and TLA edited subsequent manuscript drafts.

## Notes

https://zenodo.org/record/7750374#.ZE0uQezMLlw

### Competing Interest Statement

The authors have declared no competing interest.

