## Supplemental Information for "Plant neopolyploidy and genetic background differentiates the microbiome of duckweed across a variety of natural freshwater sources"

Table S1: Location of diploid progenitors used to create the pairs of diploid-neopolyploid genetic lineages of *Spirodela polyrhiza*.

| Genetic Lineage | Site Name | Latitude | Longitude |
| --- | --- | --- | --- |
| SP.00001 | Brady’s Run, PA | 40.73078333 | -80.35900000 |
| SP.00005 | Deer Lake, PA | 40.62100000 | -79.82675000 |
| SP.00041 | Moraine Parke, PA | 40.94098333 | -80.09125000 |
| SP.00043 | Pymatuning, OH | 41.60588333 | -80.53876667 |

Table S2: Locations of ponds used as sources of bacterial inoculum. The numbers associated with each location name correspond to the map in figure 1 of the main text. Values for pH are represented by the average ± standard error of three measurements taken from each pond source. Values of nitrate, phosphate, TDS (total dissolved solids), CaCO3 (calcium carbonate), and sulfate are in milligrams per liter.

| **Location Name** | **Latitude** | **Longitude** | **Elevation** | **pH** | **Nitrate** | **Phosphate** | **TDS** | **CaCO3** | **Sulfate** |
| --- | --- | --- | --- | --- | --- | --- | --- | --- | --- |
| **1:** Sinnemahoning State Park, PA, USA | 41.456648 | -78.058348 | 314 | 6.34 ± 0.04 | 0.0290 | 0.03407 | 52 | 26 | 6.2 |
| **2:** Elmira, NY, USA | 42.144978 | -76.91127 | 278 | 6.74 ± 0.03 | 0.0356 | 0.02996 | 700 | 155 | 22.4 |
| **3:** Stewart Park, NY, USA | 42.460074 | -76.504348 | 114 | 7.02 ± 0.07 | 0.0087 | 0.19212 | 292 | 130 | 7.9 |
| **4:** Tifft Nature Preserve, NY, USA | 42.84656 | -78.852221 | 175 | 7.03 ± 0.04 | 0.0167 | 0.05892 | 231 | 222 | 3.9 |
| **5:** Meadville, PA, USA | 41.586858 | -80.242116 | 323 | 6.66 ± 0.11 | 0.0100 | 0.042 | 146 | 131 | 4.7 |
| **6:** State Game Lands 151, PA, USA | 41.101172 | -80.128774 | 373 | 7.05 ± 0.01 | 0.0582 | 0.0192 | 148 | 79 | 3.2 |
| **7:** Virgin Run Lake, PA, USA | 40.039394 | -79.725769 | 365 | 8.59 ± 0.1 | 0.0447 | 0.02305 | 120 | 83 | 16.2 |
| **8:** Deegan Lake, WV, USA | 39.266598 | -80.248239 | 320 | 8.12 ± 0.12 | 0.0380 | 0.01525 | 122 | 56 | 14.8 |
| **9:** Cranberry Glade Lake, PA, USA | 39.907075 | -79.370431 | 702 | 5.72 ± 0.1 | 0.0100 | 0.01561 | 40 | 15 | 3 |
| **10:** Panther Hollow Lake, PA, USA | 40.436826 | -79.947263 | 271 | 6.87 ± 0.07 | 0.5806 | 0.07629 | 372 | 131 | 41.1 |

Table S3: Summary of a random forest model that classified samples based on their bacterial communities. Out of bag error rates represents the overall error rate of the model in classifying the ploidy level of the host plant, and the associated error rates with either ploidy level (2x: diploid; 4x: neopolyploid) represents the specific number of times the model misclassified the ploidy of a sample.

| **Out of bag error rate**: 26.88 % | | | |
| --- | --- | --- | --- |
|  | **2x** | **4x** | **Error** |
| **2x** | 120 | 40 | 25.00 % |
| **4x** | 46 | 114 | 28.75 % |

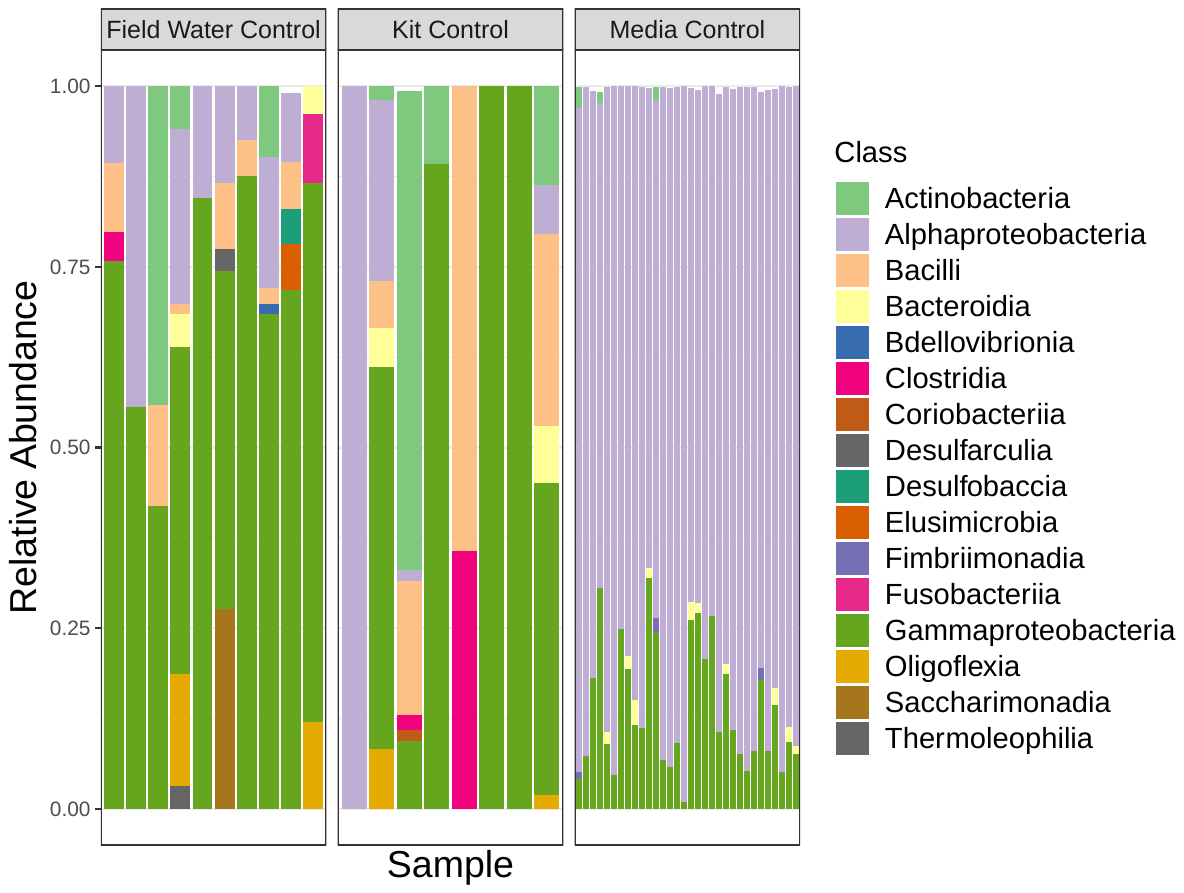

Figure S1: Relative abundance plot of amplified sequence variants (ASVs) detected in our field water controls, sequencing kit controls, or axenic *Spirodela polyrhiza* grown in sterile media controls. A full taxonomy table of these ASVs that were removed in all downstream analyses is available from Appendix S1.
